# Characterizing features affecting local ancestry inference performance in admixed populations

**DOI:** 10.1101/2024.08.26.609770

**Authors:** Jessica Honorato-Mauer, Nirav N. Shah, Adam X. Maihofer, Clement C. Zai, Sintia Belangero, Caroline M. Nievergelt, Psychiatric Genomics Consortium for PTSD Ancestry Working Group, Marcos Santoro, Elizabeth Atkinson

**Affiliations:** Department of Molecular and Human Genetics, Baylor College of Medicine, Houston, TX, 77030, USA; Department of Psychiatry, School of Medicine, University of California at San Diego, La Jolla, CA 92093, USA; Department of Psychiatry, Institute of Medical Science, Laboratory Medicine and Pathobiology, University of Toronto; Department of Morphology and Genetics, Universidade Federal de São Paulo, São Paulo, 04023-062, Brazil; Department of Biochemistry, Molecular Biology Division, Universidade Federal de São Paulo, São Paulo, 04023-062, Brazil; The Jan and Dan Duncan Neurological Research Institute, Texas Children’s Hospital, Houston, TX 77030, USA

## Abstract

In recent years, significant efforts have been made to improve methods for genomic studies of admixed populations using Local Ancestry Inference (LAI). Accurate LAI is crucial to ensure downstream analyses reflect the genetic ancestry of research participants accurately. Here, we test analytic strategies for LAI to provide guidelines for optimal accuracy, focusing on admixed populations reflective of Latin America’s primary continental ancestries – African (AFR), Amerindigenous (AMR), and European (EUR). Simulating LD-informed admixed haplotypes under a variety of 2 and 3-way admixture models, we implemented a standard LAI pipeline, testing three reference panel compositions to quantify their overall and ancestry-specific accuracy. We examined LAI miscall frequencies and true positive rates (TPR) across simulation models and continental ancestries. AMR tracts have notably reduced LAI accuracy as compared to EUR and AFR tracts in all comparisons, with TPR means for AMR ranging from 88-94%, EUR from 96-99% and AFR 98-99%. When LAI miscalls occurred, they most frequently erroneously called European ancestry in true Amerindigenous sites. Using a reference panel well-matched to the target population, even with a lower sample size, LAI produced true-positive estimates that were not statistically different from a high sample size but mismatched reference, while being more computationally efficient. While directly responsive to admixed Latin American cohort compositions, these trends are broadly useful for informing best practices for LAI across other admixed populations. Our findings reinforce the need for inclusion of more underrepresented populations in sequencing efforts to improve reference panels.

## Introduction

Admixed populations present a challenge in genome-wide analyses, as their genomes contain components from different continental ancestries, which vary from person to person along the genome, even if two people have the same overall ancestry proportions. This makes it statistically challenging to control for population structure, which can bias tests if left uncorrected^1,2^. Despite recent advances in complex trait genetics, limitations remain in our understanding of the architecture of genetic disorders in diverse populations due to their exclusion from many genomic studies^3–5^. For example, Latin American (LatAm) populations currently represent only 1.3% of all genome-wide association studies (GWAS) samples, despite accounting for 8.4% of the world population and contributing disproportionately to GWAS findings^6^. As large-scale efforts begin to focus more heavily on admixed groups, there is an unmet need for the design of well-suited pipelines to appropriately study these underrepresented populations^7^.

Local Ancestry Inference (LAI) is a machine learning approach for assigning each genomic region to a specific ancestry group by comparing phased genotype data to a reference sample containing representative phased whole genome sequence data. This method allows the researcher to assign the ancestral origin for each ancestry component for subsequent analyses, providing a better framework for control over population structure than only considering global admixture, as is accomplished with including principal components as covariates in statistical testing. There are several newly established analysis methods and tools that are tailored to admixed populations that implement LAI to deconvolute continental ancestry components in admixed samples. This work has shown that LAI can improve discovery power in genome-wide association studies for identifying ancestry-specific hits^8^, improve in polygenic risk scoring^9^, can be meaningful in evolutionary research^10^, assist in characterizing gene-gene interactions^11^, as well as provide more meaningful patient stratification in precision medicine^12,13^. To ensure and maximize the success of improving accuracy and statistical power in analyses involving LAI for admixed samples, is it paramount that local ancestry is correctly called in the individual haplotypes. However, limitations in available reference panels for many understudied populations hinder analysis, and the reference panel characteristics and algorithm parameters that result in optimal local ancestry inference accuracy across populations are still not firmly established.

One of the main features affecting LAI analysis is the reference panel used to infer local ancestry on the target sample. Reference panels are broadly required for genomic pipelines including LAI, yet are often sparse for admixed populations, particularly those who have some Amerindigenous (AMR) ancestry. Further, many reference samples are themselves admixed, which complicates assigning ancestral tracts, often resulting in admixed populations having diminished accuracy if reference panel homogeneity is assumed. As such, LAI may perform differentially for different ancestry components or populations such that there is an unmet need in establishing guidelines for best practice for diverse cohorts who may not have large, well-matched banks of reference samples to train algorithms on.

Here, we comprehensively test strategies for conducting LAI using existing reference resources with simulated “truth” genomic datasets reflective of the demographic history of Latin America to identify features that result in the best true positive rates. Latin American (LatAm) populations have a complex ancestry makeup resulting from past admixture events from multiple continental areas. Though the specific patterns vary between different geographic regions, historical admixture events generally involved substantial contributions from Amerindigenous (AMR), European (EUR) and/or African (AFR) populations^14–16^. Thus, the genomes of Latin American individuals are complex mosaics of different ancestral tracts that vary in length depending on the historical timing of when pulses of admixture occurred. We wish to highlight that we only describe inference of individuals genetic ancestry throughout this manuscript, rather than any metric of self-identification.

In our tests, we modify key parameters affecting LAI performance, including: 1) how well matched the reference panel is to the sample, 2) the absolute size of the panel, 3) the presence of admixture in the reference sample, 4) genomic data type/the number of variants (i.e. genotyping arrays vs whole genome sequencing data), as well as 5) demographic features of the cohort (global admixture proportions and timing of admixture events), and 6) parameter selection in LAI models (e.g. window size, number of EM iterations) (Figure 1). This informs best practice for researchers when conducting LAI on LatAm and other admixed populations to produce the highest accuracy results.

**Figure 1.**
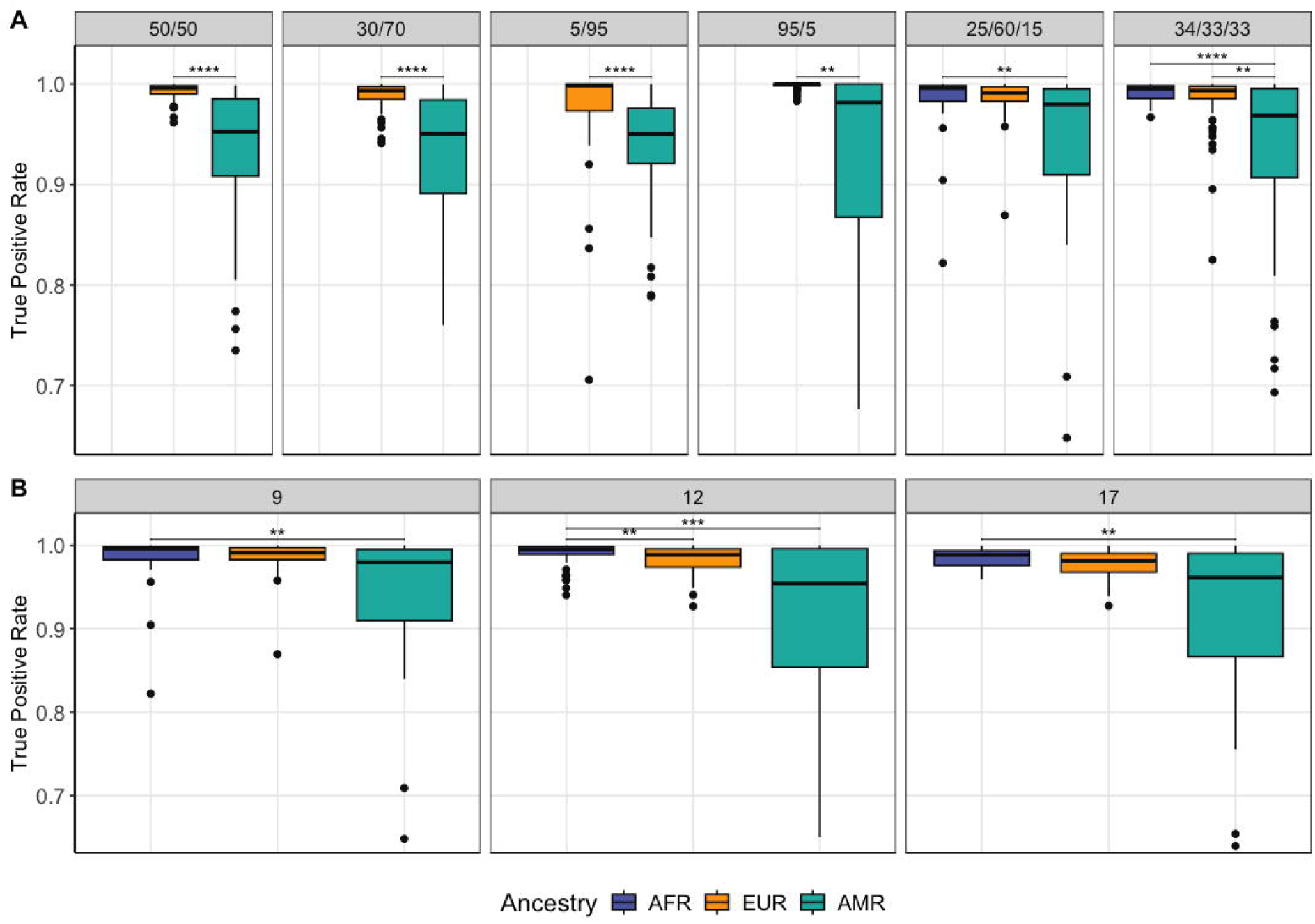
A) True positive rates for LAI in six simulated cohorts with varying proportions of 2 or 3-way admixture between AFR/EUR/AMR (displayed in order of decreasing mean TPR). These simulated haplotypes consist of chromosome 1 and considered a pulse of admixture at 9 generations ago. B) True positive rates for LAI in varying generations since admixture models for the simulated haplotype data. The haplotypes in this comparison had 15% AMR/ 60% EUR/ 25% AFR proportions of admixture in all autosomes. Significance level: * <= 0.05, ** <= 0.01, *** <= 0.001, **** <= 0.0001.

## Methods

### Dataset generation and quality control

To generate both a simulated truth dataset and comparison reference panels, we used data from the jointly called dataset of 1000 genomes (1KG) and Human Genome Diversity Project (HGDP) on GRch38^17–19^. For our three-way admixed analyses, we used data from Amerindigenous populations from HGDP (Karitiana, Surui, Colombian, Maya, Pima) as well as the Peruvians from Lima, Peru (PEL) and, in one test, East Asian (EAS) populations from 1KG to capture AMR ancestry. We wish to clarify that in this manuscript we use the term ‘AMR’ to refer to the Amerindigenous ancestry present in modern-day Latin America, rather than as a population label for admixed American samples, as has been occasionally done in prior efforts. Each of the AMR populations from HGDP was randomly split in half, with one half used for admixture simulations (N = 31) and the other used as reference sample for LAI (N=31). To keep sample sizes balanced between ancestries for simulations, we selected 30 Iberians in Spain (IBS) samples from 1KG to capture southern European ancestry and 30 Yoruba in Ibadan, Nigeria (YRI) samples from 1KG for western African ancestry. For the reference panel for LAI, we used the remaining samples from IBS and YRI populations (N=77 each) and the other half of the AMR samples, to represent a common analytic scenario. We filtered to keep only unrelated individuals and excluded multiallelic or duplicated variants, as well as those with a missingness rate > 10% and minor allele frequency < 0.5%. Genomic phasing of the complete dataset was conducted using SHAPEIT4^20^, and after phasing we subset the populations of interest as described below for our various simulations, with some samples used to model truth individuals and some used as LAI reference. By using distinct samples for our sample generation and reference panels we have an unbiased estimate of accuracy, at the cost of reducing the reference sample size.

### Simulating truth admixed haplotypes

Because of the reference sample size limitations, we simulated 60 haplotypes for each admixed cohort. Sample sizes for the reference component ancestries were selected to be equivalent to avoid biases due to unbalanced representation, and the terminal node size flag (-n 5) was implemented in LAI runs to further account for any sample size differences.

Latin America is a highly diverse region, and cohort admixture proportions vary widely depending on the country and even within each country^14,15,16^. Here, we simulated cohorts with six global ancestry patterns based on common ancestry proportions observed across Latin America. Briefly, in these simulations, one pulse of admixture is simulated at a designated point in time with specified global ancestry proportions contributed from the relevant source populations, after which haplotypes taken from the reference dataset are copied from the previous generation until the present, with tract switches informed by a recombination map. We used here the hg38 HapMap combined recombination map which includes representatives from relevant global populations^21^. This results in a simulated truth dataset that is highly similar to modern empirical LatAm cohorts but has known phase and local ancestry which can be used for method benchmarking.

We tested four two-way models of AMR/EUR admixture and two three-way models of AMR/EUR/AFR admixture. In the two-way models, we compared the effects of ancestry proportions in models with: average proportions for a two-way LatAm individual (70% AMR/30% EUR, termed ‘average two-way model’)^15^, even AMR/EUR proportions (‘even model’), and two models to analyze the effect of extreme ancestry proportions, each with 5% of one ancestry and 95% of the other (‘extreme models’). This allows us to assess the performance that may be expected in a typical two-way admixed empirical sample, as well as assess features of sample composition influencing accuracy performance.

For three-way models, we tested a model of average proportions for a three-way admixed Latin American individual (15% AMR/60% EUR/25% AFR-average proportions for a Brazilian individual, termed ‘average 3-way model’)^22^ and an even-proportioned model. Simulations were conducted using the admix-simu tool^23^. We simulated the average 3-way model in three different admixture demographic scenarios, considering a single pulse of admixture at 9 generations, 12, or 17 generations ago^14^. This allowed us to evaluate the impact of varying tract lengths on true positive LAI rates. All other models were simulated considering a single pulse of admixture at 9 generations ago for the sake of comparability. In the simulation of the admixture model that has 3-way average LatAm proportions and a pulse of admixture 12 generations ago, we used data for all autosomes to obtain the highest precision. This admixture model was landed upon as it is reflective of the intermediate admixture pulse in a population migration model for the Brazilian population according to Kehdy et al., 2015^14^. For all other simulations, we simulated only chromosome 1 for the sake of computational efficiency.

For comparisons of DNA data generation type, we created a pseudo-genotype array dataset by selecting all SNVs present in the Global Screening Array (Illumina GSA) from our WGS-density simulation reference dataset, a genotyping array that has been regularly used for non-European datasets. To test the effect of imputation on LAI accuracy, given that imputation is a typical step in cohort data processing for genomic analyses such as GWAS, we imputed the simulated haplotypes with SNP array-density sites using the TOPMed panel^24^ imputation server and filtered imputed sites with > 0.8 INFO score and MAF > 0.005.

### Local Ancestry Inference

Local Ancestry was deconvoluted using RFMix v1.5.4^25^. We used the TrioPhased option, with a base window size of 0.2 cM, terminal node size of 5, 2 Expectation–maximization (EM) iterations, with reference panels reanalyzed in EM to account for any admixture present in the reference (flags -w 0.2, -n 5, -e 2, and --use-reference-panels-in-EM, respectively), and the number of generations since admixture was specified depending on the simulation model (9, 12 or 17). For reference panel testing, we used three different reference panel combinations from HGDP and 1KG (AMR/EUR or AMR/EUR/AFR) that varied only in the AMR reference samples, given that this group has much less representation in reference panels relative to the other two ancestries. This was done to benchmark how variations in the reference for this ancestry impacts LAI accuracy with particular attention to improving AMR accuracy given the limitations of available reference resources. The three reference panels for LAI were constructed using, for EUR and AFR components respectively, the remaining IBS and YRI samples from 1KG not used in the simulations (N IBS = 77, N YRI = 77). For the AMR component, the three panels varied as follows: 1) *Well matched to the target but low sample size*: using the other half of HGDP-AMR samples (N = 30); 2) *Medium sample size containing some admixture in AMR component*: using the 1KG sample from Lima, Peru (PEL) (N = 85); 3) *Large sample size but AMR component poorly matched to the target*: 1KG-PEL (N = 85) plus 1KG East Asian (EAS) populations (N = 505). We included the EAS population on panel 3 to capture highly diverged AMR ancestry considering the demographic history of human migrations, since the ancestors of modern Amerindigenous peoples of the Americas migrated from East Asia across the Bering Strait around fifteen thousand years ago^26^. As such, AMR and EAS ancestry are less diverged than other ancestral components, but even so, this composition makes for a poorly matched reference panel to the target data. Panel 2 represents the procedure most commonly conducted in current studies. We used the LAI reference panel containing the AMR samples described in panel 1 for all comparisons that did not involve reference panel testing.

### Statistical Analysis

We quantified the LAI true positive rates (TPR) for each run to assess their respective performance. We define TPR as follows: for each run we calculated the sums of genomic positions for which a given ancestry was correctly called compared to the simulated true ancestry for that position, divided by the total number of positions for that ancestry overall in the cohort, in each simulated haplotype (Supplementary Figure 1). We analyzed the best-guess ancestry calls output by RFMix, regardless of confidence (Forward-Backward) estimates. We computed ancestry-specific TPR to assess if there was differential LAI performance depending on the background truth ancestry and tested for statistically significant differences using the Wilcoxon rank-sum test. Significance was considered when the Bonferroni-adjusted p-value (p-adj) < 0.05.

## Results

### Impact of demography and ancestry proportions on LAI performance

We compared the effect of different demographic models on LAI performance. Specifically, we assessed the impact of varying component ancestry proportions in two and three-way models as well as different generation times since an admixture pulse occurred (considering a single pulse) by simulating 9, 12 and 17 generations since admixture.

In general, LAI accuracy for a given ancestry increased as that global ancestry proportion increased (Figure 1, Table 1). A low global ancestry percentage tended both to decrease the accuracy and result in larger standard deviations, as observed in the “extreme proportions” simulations (95% EUR/5% AMR and 5% EUR/95% AMR, Figure 1). In all tested models, we additionally observed significantly lower true positive rates for the AMR component (p-adj < 0.05). This result was consistent for all chromosomes and demographic models tested (Figure 1, Figure 2, Supplementary Table S1).

**Table 1.**
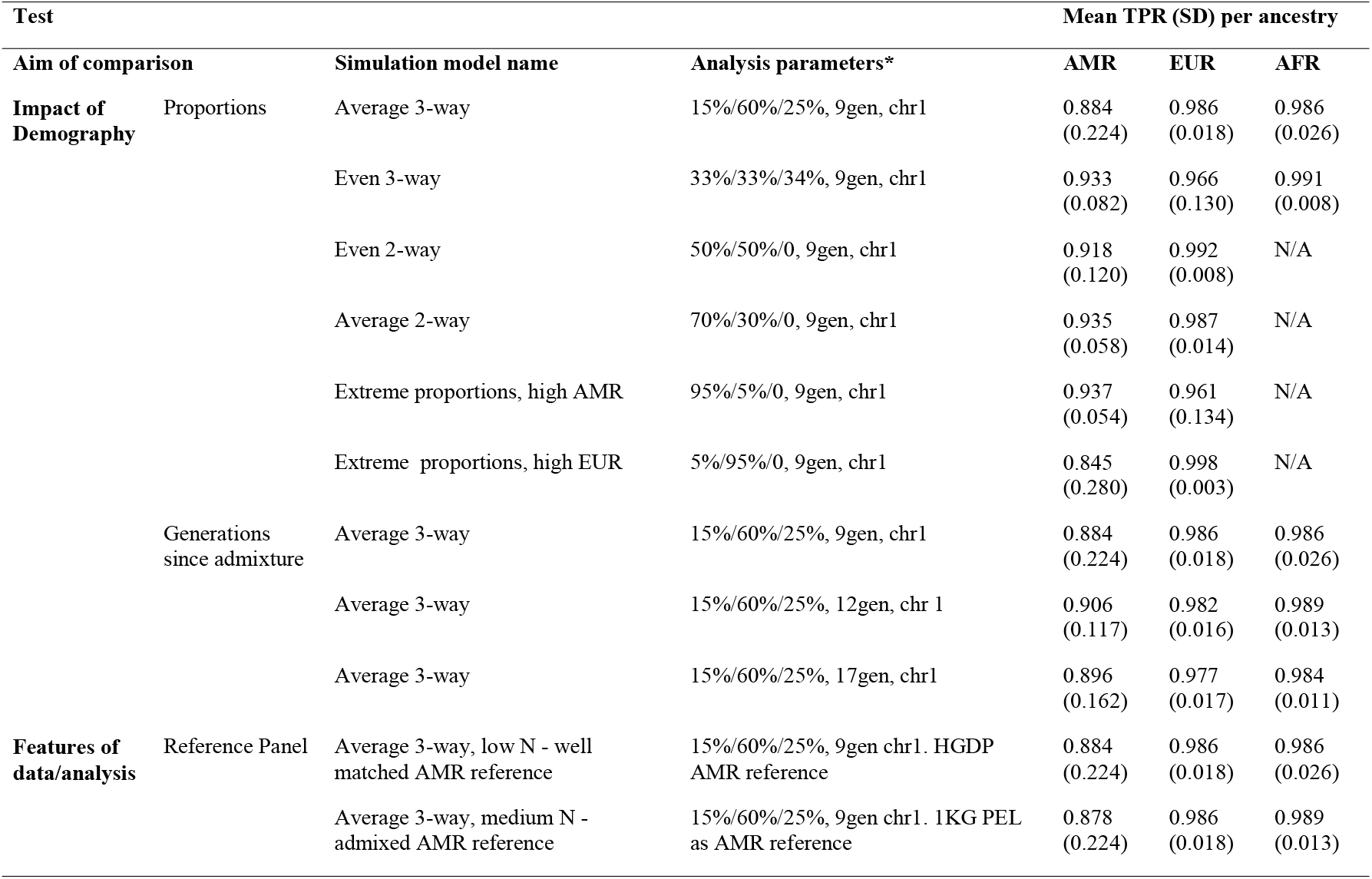
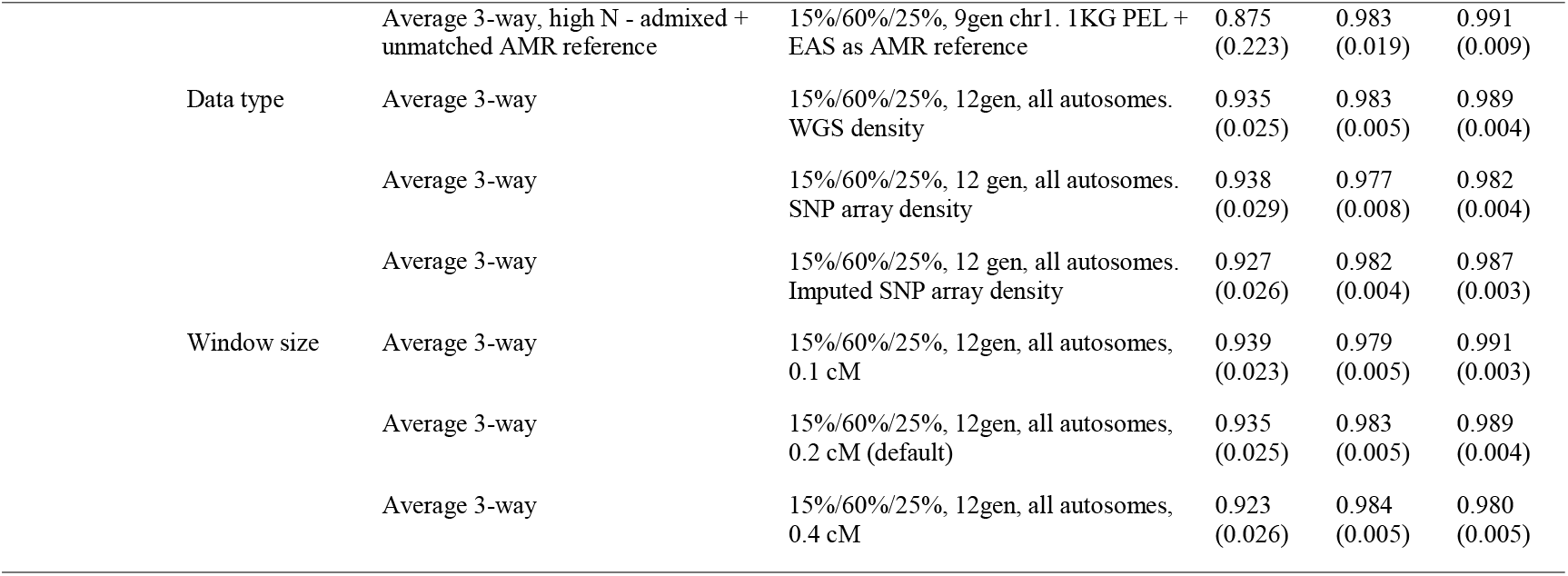
LAI true positive rate estimates per ancestry per comparison *Analysis parameters: Admixture proportions of simulated cohort (AMR/EUR/AFR), number of generations since admixture, simulated chromosome, other parameters.

| Test |  |  |  | Mean TPR (SD) per ancestry |  |  |
| --- | --- | --- | --- | --- | --- | --- |
| Aim of comparison |  | Simulation model name | Analysis parameters* | AMR | EUR | AFR |
| **Impact of Demography** | Proportions | Average 3-way | 15%/60%/25%, 9gen, chr1 | 0.884<br>(0.224) | 0.986<br>(0.018) | 0.986<br>(0.026) |
|  |  | Even 3-way | 33%/33%/34%, 9gen, chr1 | 0.933<br>(0.082) | 0.966<br>(0.130) | 0.991<br>(0.008) |
|  |  | Even 2-way | 50%/50%/0, 9gen, chr1 | 0.918<br>(0.120) | 0.992<br>(0.008) | N/A |
|  |  | Average 2-way | 70%/30%/0, 9gen, chr1 | 0.935<br>(0.058) | 0.987<br>(0.014) | N/A |
|  |  | Extreme proportions, high AMR | 95%/5%/0, 9gen, chr1 | 0.937<br>(0.054) | 0.961<br>(0.134) | N/A |
|  |  | Extreme proportions, high EUR | 5%/95%/0, 9gen, chr1 | 0.845<br>(0.280) | 0.998<br>(0.003) | N/A |
|  | Generations since admixture | Average 3-way | 15%/60%/25%, 9gen, chr1 | 0.884<br>(0.224) | 0.986<br>(0.018) | 0.986<br>(0.026) |
|  |  | Average 3-way | 15%/60%/25%, 12gen, chr 1 | 0.906<br>(0.117) | 0.982<br>(0.016) | 0.989<br>(0.013) |
|  |  | Average 3-way | 15%/60%/25%, 17gen, chr1 | 0.896<br>(0.162) | 0.977<br>(0.017) | 0.984<br>(0.011) |
| **Features of data/analysis** | Reference Panel | Average 3-way, low N - well matched AMR reference | 15%/60%/25%, 9gen chr1. HGDP AMR reference | 0.884<br>(0.224) | 0.986<br>(0.018) | 0.986<br>(0.026) |
|  |  | Average 3-way, medium N - admixed AMR reference | 15%/60%/25%, 9gen chr1. 1KG PEL as AMR reference | 0.878<br>(0.224) | 0.986<br>(0.018) | 0.989<br>(0.013) |

**Table 1 Legend:** \*Analysis parameters: Admixture proportions of simulated cohort (AMR/EUR/AFR), number of generations since admixture, simulated 571 chromosome, other parameters.
|  |  |  |  |  |  |
| --- | --- | --- | --- | --- | --- |
|  | Average 3-way, high N - admixed + unmatched AMR reference | 15%/60%/25%, 9gen chr1. 1KG PEL + EAS as AMR reference | 0.875<br>(0.223) | 0.983<br>(0.019) | 0.991<br>(0.009) |
| Data type | Average 3-way | 15%/60%/25%, 12gen, all autosomes. WGS density | 0.935<br>(0.025) | 0.983<br>(0.005) | 0.989<br>(0.004) |
|  | Average 3-way | 15%/60%/25%, 12 gen, all autosomes. SNP array density | 0.938<br>(0.029) | 0.977<br>(0.008) | 0.982<br>(0.004) |
|  | Average 3-way | 15%/60%/25%, 12 gen, all autosomes. Imputed SNP array density | 0.927<br>(0.026) | 0.982<br>(0.004) | 0.987<br>(0.003) |
| Window size | Average 3-way | 15%/60%/25%, 12gen, all autosomes, 0.1 cM | 0.939<br>(0.023) | 0.979<br>(0.005) | 0.991<br>(0.003) |
|  | Average 3-way | 15%/60%/25%, 12gen, all autosomes, 0.2 cM (default) | 0.935<br>(0.025) | 0.983<br>(0.005) | 0.989<br>(0.004) |
|  | Average 3-way | 15%/60%/25%, 12gen, all autosomes, 0.4 cM | 0.923<br>(0.026) | 0.984<br>(0.005) | 0.980<br>(0.005) |

**Figure 2.**
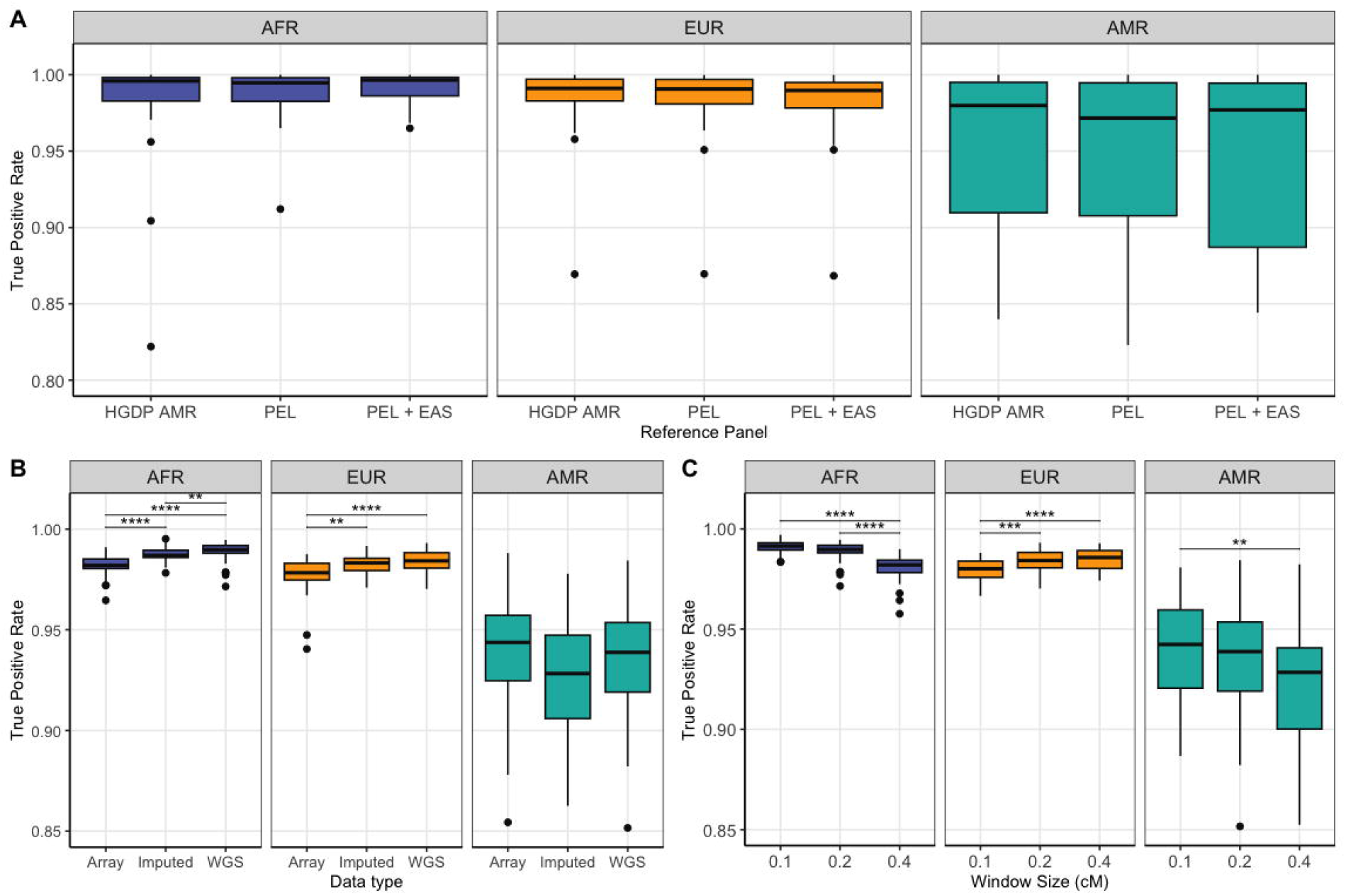
A) True positive rates for LAI in three reference panel comparisons that vary in the AMR component, separated by ancestry component. Benchmarking was run on the model reflecting a pulse of admixture at 9 generations ago with 15% AMR/ 60% EUR/ 25% AFR proportions in chromosome B) True positive rates for LAI in WGS vs. SNP array (GSA) vs. Imputed data. Benchmarking was run on the model reflecting a pulse of admixture at 12 generations ago with 15% AMR/ 60% EUR/ 25% AFR proportions in all autosomes. C) True positive rates for LAI runs varying the RFMix window size parameter in centimorgans (cM). Benchmarking was run on the model reflecting a pulse of admixture at 12 generations ago with 15% AMR/ 60% EUR/ 25% AFR proportions in all autosomes. Significance level: * <= 0.05, ** <= 0.01, *** <= 0.001, **** <= 0.0001.

The number of generations since admixture had a slight impact on TPR, which modestly decreased in older admixture events (17 generations ago) compared to more recent ones (9 generations ago). We observed statistically significant differences in TPR between the 9 and 12 generations models in the AFR and EUR components, and 12 and 17 generations models in the AFR component (p-adj < 0.05). In all models the 17 generations since admixture model had the lowest accuracy (Figure 1-B, Table 1, Supplementary Table S1). This is likely due to increased difficulty in painting short ancestry tracts; the further back in time a pulse occurred, the shorter the relative ancestral tracts will be in the current day as recombination breaks ancestral stretches down over time^27^. The general trends observed in relative TPR per ancestry proportion were the same regardless of admixture pulse generation times.

### Impact of reference panel compositions

For benchmarking the performance of different reference panel compositions, we tested multiple AMR/EUR and AMR/EUR/AFR reference panel combinations comprising individuals from the Human Genome Diversity Project (HGDP) and the Thousand Genomes Project (1KG). We organized our test reference panels to reflect: 1) a very well-matched panel but with low sample size; 2) a moderate sized panel that includes admixed individuals in the reference versus restricting to only homogeneous individuals; or 3) a very large reference but that is poorly matched.

The average TPRs were similar across the three tested reference panels, and had no statistically significant differences (p-adj <0.05, Figure 2-A, Supplementary table S2), however the small but well-matched reference panel (N=184) resulted in considerably faster running time compared to the admixed AMR reference panel (N=239) and the large but unmatched reference panel (N=659), which took three and sixteen times longer to complete a LAI run for chromosome 1, respectively. Figure 2-A and Table 1 summarize the TPR results for each reference panel.

### Impact of genetic data type

We assessed the impact of genetic data type by comparing true positive rates (TPR) of LAI with simulations generated from WGS-density reference, a subset of SNP array variants, and a dataset of imputed variants (using the subset of SNP array variants as input), in the average 3-way (15% AMR/ 60% EUR/ 25% AFR) admixture model simulation considering 12 generations since admixture. We selected genomic variants targeted by the GSA chip for these tests, as this is a commonly used array for diverse datasets.

LAI run on the WGS-density simulated dataset achieved better TPR for AFR and EUR ancestry components than the SNP array-density dataset (p-adj < 0.05), likely due to fuller haplotype coverage, but was roughly 6 times slower to complete for all autosomes. Imputation slightly improved LAI calls for these ancestry components compared to the SNP-array only runs, indicating increased SNP density improved performance. These trends were different in the AMR component, however, in which we observed no statistically significant differences between either WGS, SNP array and imputed datasets, and observed a decrease in TPR following imputation (the lowest TPR in this component), although not statistically significant (Table 1, Figure 2-B, Supplementary table S2). Additionally, we performed a validation analysis of the imputation accuracy results by selecting only the original SNP-array sites from the LAI results of the imputed dataset, to observe if imputation changed LAI on these sites which could lead to changes in accuracy performance. We observed no significant changes in TPR compared to the full imputed dataset (Supplementary Figure 2).

### RFmix window size parameter changes do not improve LAI accuracy from the default value

As RFmix calls LA with a sliding window approach, we tested whether halving or doubling the default window size improved calls, which could change results especially at the borders of chromosomes that may only have anchoring haplotype information on one side of the window.

We found that halving the default window size to 0.1cM did not significantly change TPR for the AFR and AMR components, and significantly lowered TPR in the EUR component compared to the default 0.2 cM (p-adj < 0.0001). Doubling the default window size to 0.4 cM significantly decreased TPR for the AFR component (p-adj < 1e-5) but did not significantly change for the other components compared to the default (Figure 2-C, Table 1, Supplementary table S2). As such, we recommend retaining the default 0.2cM window size for RFmix runs.

We additionally examined the ForwardBackward probability estimates from RFMix to check how confident the algorithm estimated the wrong calls, as such confidence estimates could be a readily implemented filter to remove poorly called loci. We observed, however, that miscalls had high confidence estimates, therefore setting a stringent filter for ForwardBackward probabilities in an attempt to reduce LAI miscalls would not be sufficient to improve results.

### LAI miscalls are more frequent in certain genomic locations, but vary between cohorts

When miscalls in LAI occurred, we observed that although they may occur at any point in the genome, they were more frequent around telomere and centromere regions (Figure 3-A, 3-B, Supplementary figures 3-8). Considering sites with over 10% miscalls in both three-way model runs and considering a window of 1kb upstream and downstream, we observed that these regions can span or flank genes (Supplementary tables S3-S4), most of which have been previously associated in GWAS studies according to GWAS Catalog (Supplementary Tables S5-S6). We compared these sites with low complexity regions from the UCSC RepeatBrowser hg38 dataset and observed an overlap of >98% in both models. It is important to note, however that the sites/regions with over 10% miscalls varied between the admixture models in which we ran this analysis, therefore we do not supply a list of regions that will be more frequently miscalled, as this may vary between cohorts.

**Figure 3.**
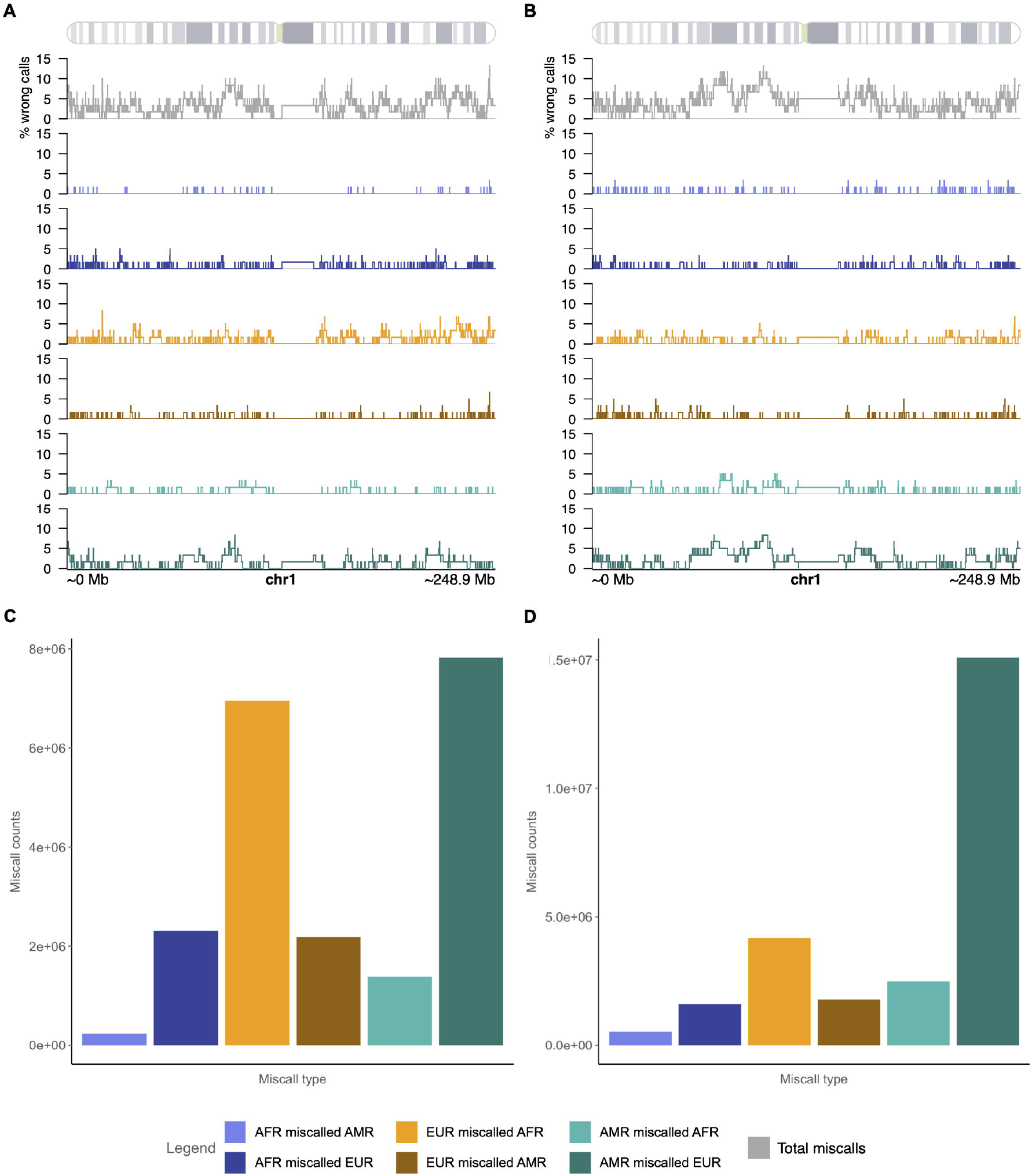
A) Percentage of wrong calls per site on chromosome 1, total and separated by error mode for LAI ran on the model reflecting a pulse of admixture at 12 generations ago with 15% AMR/ 60% EUR/ 25% AFR proportions. B) Percentage of wrong calls per site on chromosome 1, total and separated by error mode for LAI ran on the model reflecting a pulse of admixture at 12 generations ago with 33% AMR/ 33% EUR/ 34% AFR proportions. C) Miscall counts separated by error mode summing all autosomes for LAI ran on the model reflecting a pulse of admixture at 12 generations ago with 15% AMR/ 60% EUR/ 25% AFR proportions. D) Miscall counts separated by error mode summing all autosomes for LAI ran on the model reflecting a pulse of admixture at 12 generations ago with 33% AMR/ 33% EUR/ 34% AFR proportions.

### LAI miscalls occur with a consistent error mode

We summed miscall counts and divided them between “error modes”, i.e.: how many truth sites of one ancestry are being miscalled as each of the other ancestries. This allowed us to characterize trends in miscall directions and observe whether one ancestry was systematically being over-or under-called than other. We observed that the most common direction for miscalls to occur was for truth AMR sites to be incorrectly called EUR (Figure 3-C, 3-D, Supplementary figures 9-12). The second most frequent error mode was EUR positions being miscalled AFR.

## Discussion

In this study, we evaluated characteristics impacting the performance of LAI for a range of two and three-way admixed demographic models reflective of many Latin American populations. Specifically, we assessed the impact of reference panel composition, demographic features such as the proportions of major ancestry groups and number of generations since admixture in the cohort, genetic data technology (genotyping arrays versus whole genome sequencing), the impact of imputation, and LAI analytic thresholds in affecting performance for diverse cohorts. Given the high LAI accuracy observed in the literature for 2-way admixed AFR/EUR cohorts^8^, we focused our analyses in this manuscript on determining the best practices for cohorts involving AMR, as the smaller divergence time between EUR and AMR tracts poses a challenge for deconvolution, as does the particularly limited availability relevant reference individuals for AMR ancestry. Thus, we focused the construction of our reference panel tests in the service of optimizing AMR accuracy.

These benchmarks allow us to provide a set of recommendations for parameter and panel selection to achieve optimal LAI performance in LatAm populations. Specifically, comparing the performance of different reference panel compositions, we observed that there was not significant difference in accuracy across the three panels (well-matched but small sample size, medium size with some degree of admixture in the reference, and large but poorly matched to the target cohort), although we do observe a large difference in runtime, with the small but well-matched panel running substantially faster than the other panels. Given the high computational burden required by LAI, having a quicker runtime for analysis is an important point of consideration in practical use. As such, a curated reference panel reflective of the ancestries present in the target cohort appears to be the best option for LAI reference panel construction. Importantly, across all demographic and reference panel models tested, Amerindigenous ancestry tracts suffer from notably reduced accuracy as compared to European and African tracts. This is likely due to there being less representative (and less homogeneous) reference data for AMR ancestry in existing reference resources. Moving forward, it will be vital for efforts to focus on ethical recruitment of more diverse and geographically distributed reference samples in large scale data collection efforts to maximize the performance of LAI across all ancestry backgrounds.

Regarding ancestry proportions in the admixed simulations, we observed that overall, having a higher proportion of an ancestry in the simulation improved the true positive rates for that ancestry in LAI. When ancestries represent a very small (e.g. 5%) global proportion, all ancestries suffer, with AMR suffering the most.

We investigated LAI miscalls in the realistic three-way simulation models to evaluate the typical error mode when wrong calls are produced. Specifically, we examined the rates of miscalls for each ancestry component to: characterize trends in the relative amount and direction of miscalls, see if a particular ancestry was being systematically over or under-called, document if there were genomic regions where miscalls were most frequent, to identify other factors that could be driving error modes, and to assess if alterations to RFMix parameters could improve miscall rates. Investigating the typical error modes when LAI miscalls occur, we observed a much higher frequency of miscalls in the direction of calling simulated AMR regions as EUR compared to other miscall directions. This may be explained by the smaller genetic divergence between AMR and EUR haplotype tracts than either is to AFR, resulting in closer haplotype similarity. This finding implies that reference panels will need to be grown substantially to confidently assess within-continental ancestral components for many geographic regions. The specific direction of AMR/EUR miscalls being dominated in the direction of AMR to EUR rather than vice versa can be explained by the substantially (2x) larger sample size available for EUR compared to AMR. Another point of consideration is that the AMR reference samples themselves have some degree of admixture with European ancestry, which adds uncertainty to the model, though we did implement EM procedures to attempt to correct for this. These results are consistent with miscall trends observed in other studies of diverse populations^28^. As this prior cited work was done with older LAI software than RFMix, we have confirmed that this error mode is consistent between different LAI algorithms and therefore likely to be driven by the genetic data, rather than a feature specific to RFMix.

Beyond error modes, we observe that miscall regions do not appear randomly across the genome, but are most likely to fall in areas that mark the edges of haplotypes, like centromere and telomeres (Figure 3A, 3-B, Supplementary Figures 3-8). We note several areas that had elevated miscall rates (higher than 10% miscalls). ForwardBackward probabilities for the LAI algorithm were still confident in such areas and tweaking RFmix parameters was insufficient to correct them. As these regions may vary between cohorts given that different models resulted in different regions with elevated false positive rates, we do not recommend blanket masking of those observed in this study. This highlights the importance of ensuring good LAI accuracy for gene discovery and other statistical genomics efforts, as misclassification both soaks up power in LAI-informed GWAS as well as can lead to false positive associations due to technical ancestry miscalls^8^. Importantly, miscall regions may contain genes of interest, so care should be taken to validate, for example, GWAS hits in border haplotype areas that show elevated miscall rates. Inflation of miscalls at particular regions could also impact the interpretation of other statistical genetics efforts, such as admixture mapping or evolutionary scans of selection that utilize local ancestry enrichment. We observed an inflation of miscalls in low complexity regions, such as short and long interspersed nuclear elements (SINE/LINE), DNA repeats and micro-satellites, therefore additional care should be considered when analyzing these regions and/or genes in close proximity. This could be due to the fact that low complexity regions are usually more challenging to map^29,30^, and/or are evolutionarily conserved^31,32^, with little variation across ancestry groups, and therefore these regions would be more prone to error in LAI. The development of methods that incorporate repeat polymorphisms, multi-allelic variants and other complex forms of genetic variation in genome-wide analyses may help improve LAI accuracy.

Examining how different DNA data types impact LAI performance, we observe that WGS and SNP array simulated data resulted in similar TPR estimates for genotyped sites, although having more variants in the dataset improved estimates. We also observe that LAI performs nearly as well on imputed data as directly genotyped data when a large and diverse reference panel is used. This suggests that, provided imputation can be performed with a representative reference panel, LAI calls on imputed data may be confidently utilized for downstream efforts. Out of an abundance of caution, we recommend setting a stringent INFO threshold (e.g. 0.8) for imputed sites to ensure high confidence calls. We note, additionally, that non-significant differences in TPR in the context of this work do not mean that the differences that we observe are not relevant since small differences in LAI accuracy can impact statistical power in downstream applications^8^ and may represent a difference observed in a large number of sites in the genome. Expanding available reference samples to contain representative haplotypes from diverse and understudied populations would improve the quality of imputation as well as LAI.

Of course, this work has some important limitations which must be considered. As the focus of the present study is LatAm populations, we limited our demographic models to those involving 2-way admixture between AMR and EUR or 3-way admixture between AMR, EUR, and AFR, which represents the majority of Latinx populations. We note, however, that some LatAm populations have other patterns than those directly benchmarked here. Despite this, the broader trends in LAI performance identified in this work should hold across demographic models beyond the specific use cases simulated in this manuscript. We also note that while we appreciate that there is a high level of diversity within continental regions^14,33^, only continental-level ancestry was able to be assessed here due to limitations in available reference panel geographic coverage. Similarly, having a small number of available reference AMR samples limited the number of individuals available for simulating and running LAI, which limits variability in the data for this component in comparison to EUR and AFR. Improved LAI call rates and finer scale LAI resolution would be possible in the future if reference panels are expanded. Regarding software, we have benchmarked only RFMix v1 in this work, as prior work has demonstrated that RFMixv1 performed the best in comparison to other methods for multi-way admixed samples^34^. We expect the trends observed here to be consistent across LAI software, though further benchmarking would be needed to confirm this.

In conclusion, in our reference panel benchmarking, the best cost-benefit in terms of LAI accuracy and speed is to use a well-matched reference even if it has a lower sample size. Examining the ancestry-specific performance of LAI across reference panels, we observed consistently lower performance for the AMR ancestry component across all simulation settings compared to EUR and AFR. Unfortunately, this inequity could not be overcome by any of the tested modifications to reference panel, LAI software parameters, or features of genetic data. The best way to improve AMR performance would be to increase the well-matched reference panel’s sample size, underscoring the importance of furthering recruitment of larger and more representative reference samples for understudied populations. Given the high proportion of the global population that contains admixed ancestry and the fact that populations are getting increasingly admixed over time^35^, it is timely to establish the optimal methods for well-calibrated genomic analyses in admixed populations.

## Acknowledgments

This study was supported by Fundação de Amparo à Pesquisa do Estado de São Paulo (FAPESP; fellowship number: 2021/09584-1). Financial support for the PTSD-PGC Ancestry Working Group was provided by the National Institute of Mental Health (NIMH; R01MH106595). EGA was supported by the National Institute of Mental Health (K01 MH121659, R01 HG012869), the Caroline Wiess Law Fund for Research in Molecular Medicine, and the ARCO Foundation Young Teacher-Investigator Fund at Baylor College of Medicine.

## Author Contributions

J.M. conducted analysis and wrote the manuscript. A.X.M. and N.N.S. assisted with software. C.Z. reviewed the manuscript. C.M.N. and S.B. advised on the project. E.G.A. and M.S. conceptualized, supervised, and funded the project as well as contributed to the manuscript. All authors reviewed and approved the final manuscript.

## Web Resources

Admix-simu: https://github.com/williamslab/admix-simu/

RFMix V.1 https://github.com/indraniel/rfmix

Shapeit v4 https://odelaneau.github.io/shapeit4/

Pipeline used to prepare data for RFMix v1: https://github.com/armartin/ancestry_pipeline/

R package utilized to create TPR figures: https://phanstiellab.github.io/plotgardener/

UCSC RepeatBrowser: https://repeatbrowser.ucsc.edu/data/

TOPMed Imputation Server: https://imputation.biodatacatalyst.nhlbi.nih.gov/

Thousand Genomes Project: http://ftp.1000genomes.ebi.ac.uk/.

The Human Genome Diversity Project: ftp://ngs.sanger.ac.uk/production/hgdp/hgdp_wgs.20190516/statphase/.

Jointly called HGDP+1kG: https://gnomad.broadinstitute.org/downloads#v3-hgdp-1kg

HapMap GRCh38 recombination map: http://bochet.gcc.biostat.washington.edu/beagle/genetic_maps/plink.GRCh38.map.zip

## Data/Code Availability

Code generated in this project for simulating admixed data and quantifying LAI true positive rates is freely available on github at https://github.com/Atkinson-Lab/LAI-sims-accuracy.

