## Supplemental Figures for "Characterizing features affecting local ancestry inference performance in admixed populations"

**Supplementary Figures**

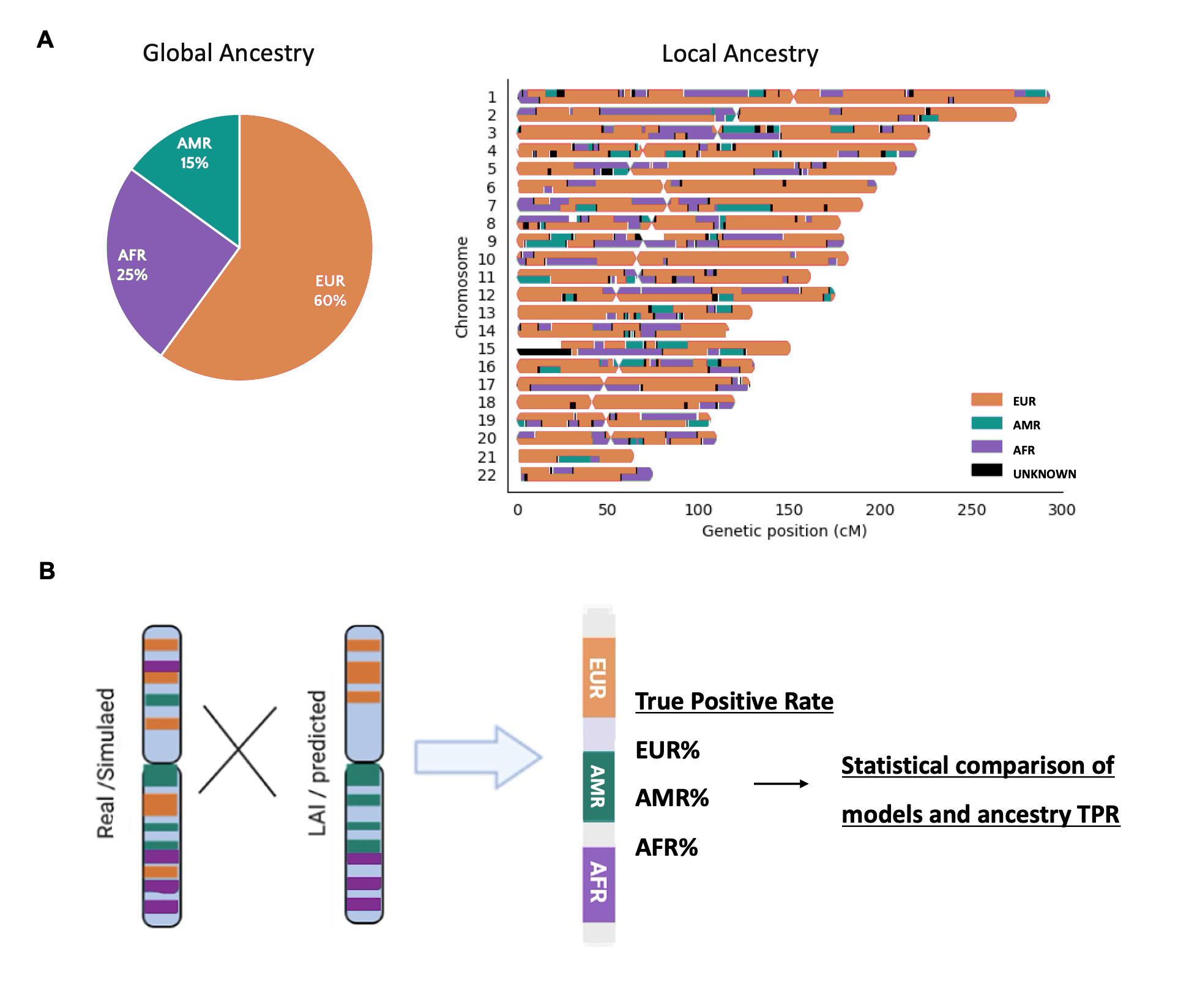

**Supplementary Figure 1:** Study design. A) Admixed populations' genomes have contributions from multiple populations. A single admixed individual has different local ancestry patterns from others in the same population, even if their global ancestry proportions are the same. This figure illustrates an example of a three-way (EUR/AMR/AFR) admixed individual. B) We simulated admixed haplotypes under a variety of admixture proportions to obtain a "truth" local ancestry dataset and ran LAI on these haplotypes to evaluate the LAI runs' true positive rates also under a variety of analytic scenarios for 2-way and 3-way admixtures: even, average and extreme ancestry proportions, different generations since admixture, levels of matching and different sizes of the reference, WGS vs imputed SNP array vs unimputed SNP array, and different window sizes for ancestry inference.

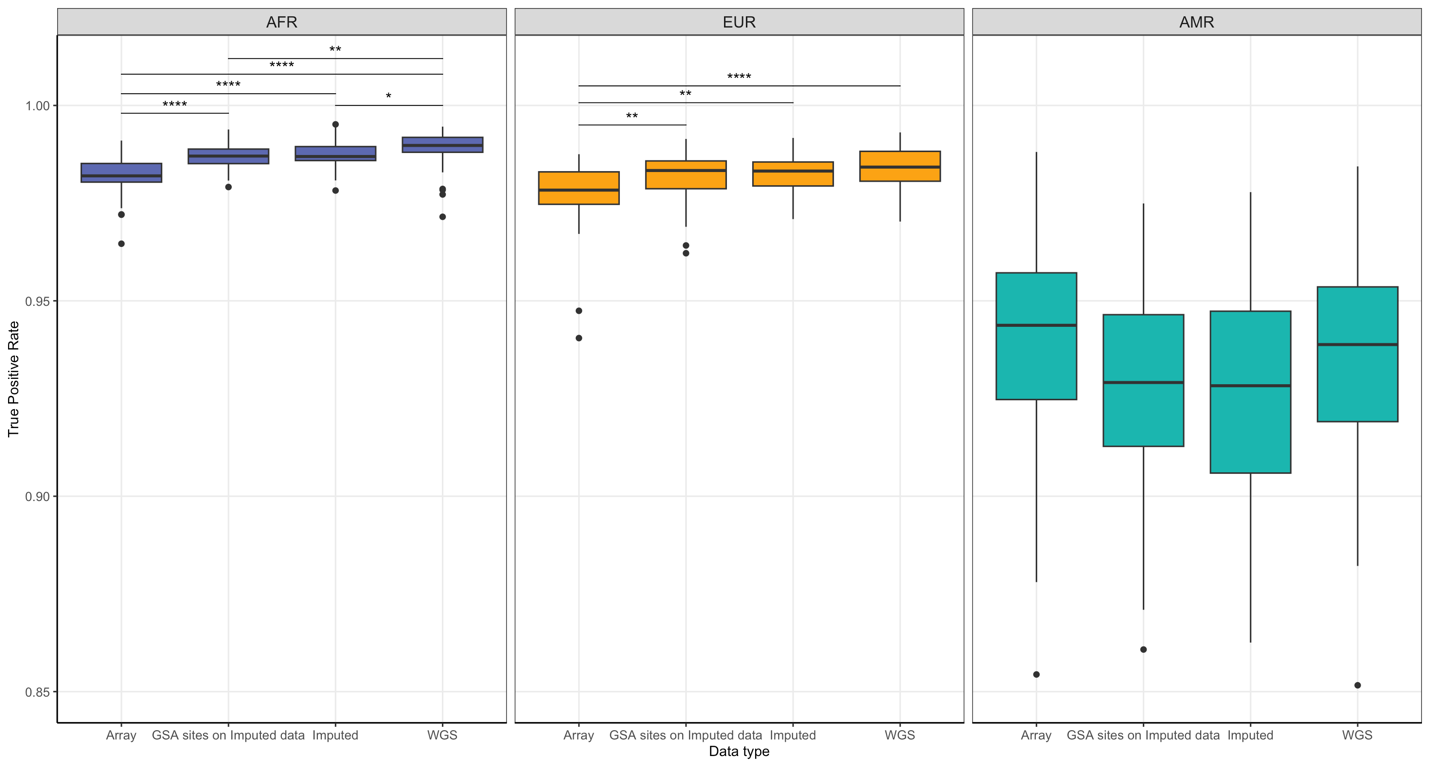

**Supplementary Figure 2:** Evaluating the impact of imputation on LAI TPR. In the "GSA sites on Imputed data" group, only original array sites were considered for TPR calculation after running LAI on imputed data.

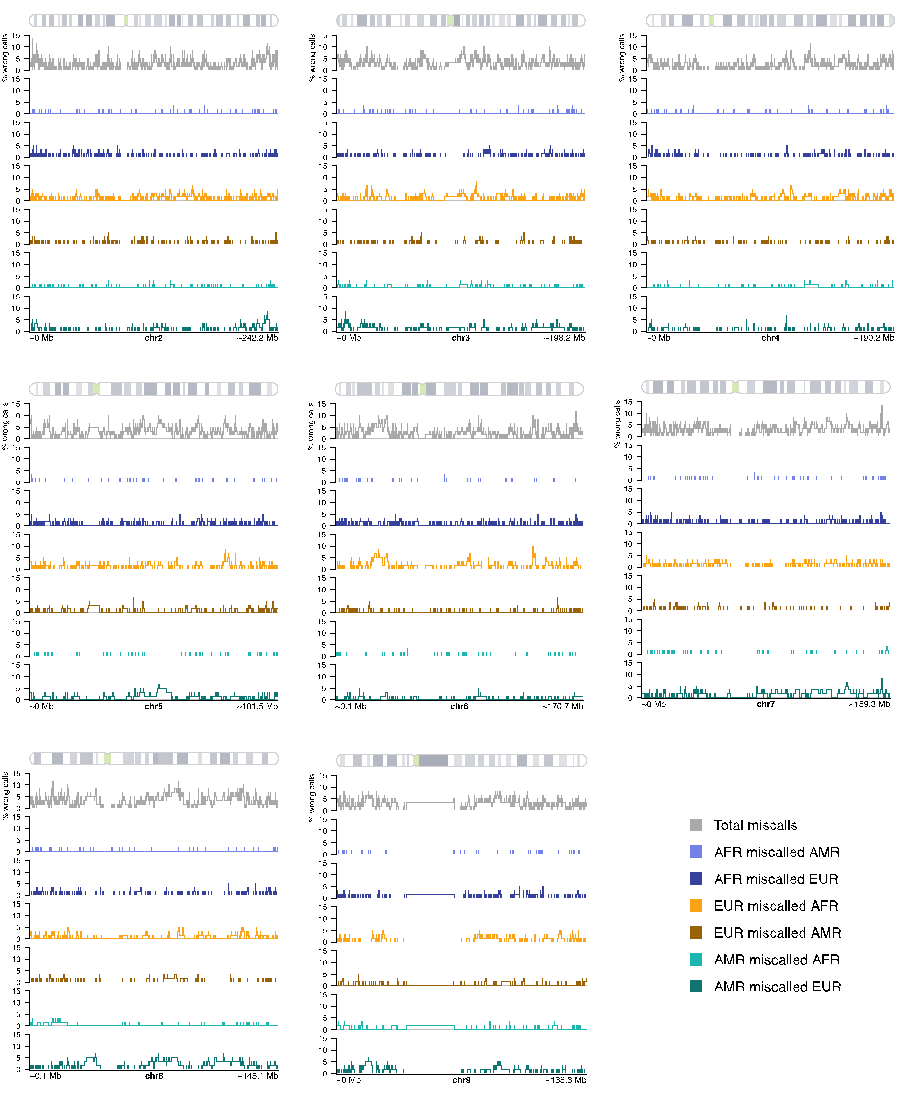

**Supplementary Figure 3:** Percentage of wrong calls per site on chromosomes 2-9 The total is shown in gray, as well as for each error mode for LAI. Results are from the ‘average 3-way’ model reflecting a pulse of admixture at 12 generations ago with 15% AMR/ 60% EUR/ 25% AFR proportions.

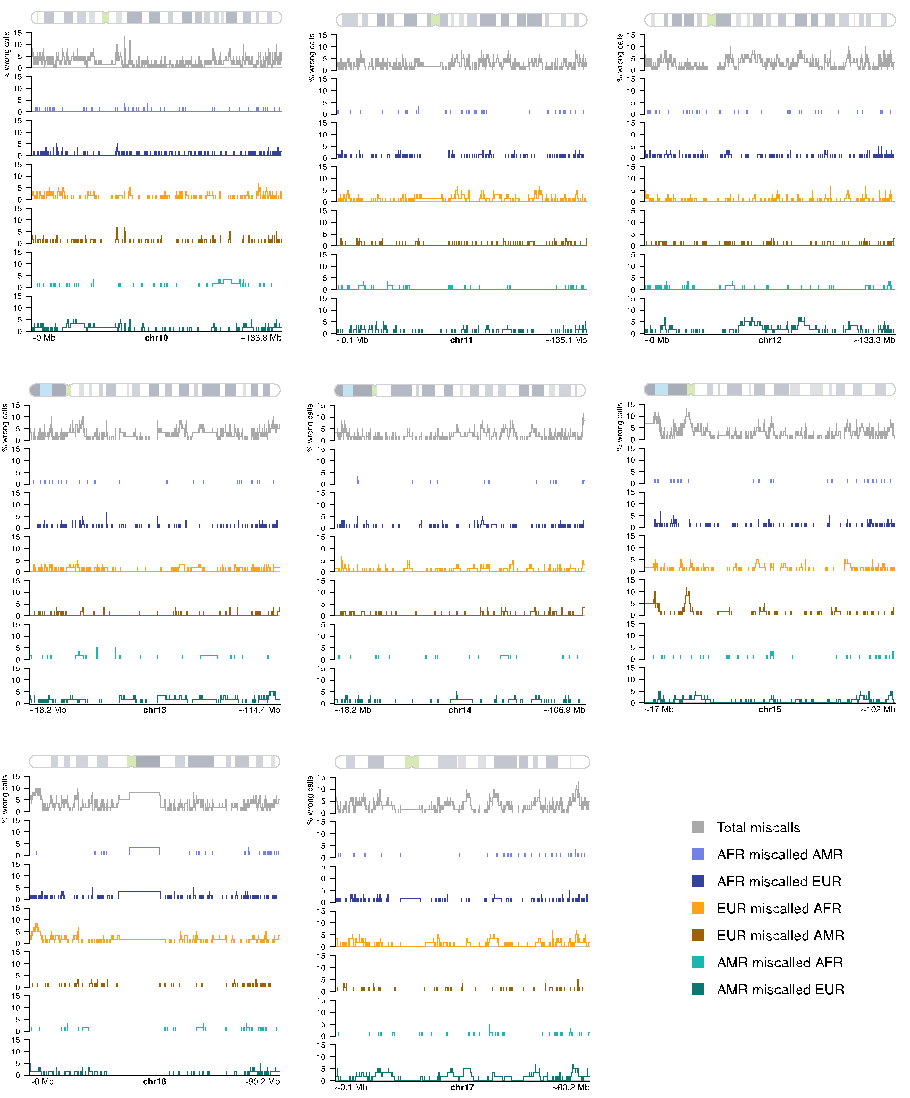

**Supplementary Figure 4:** Percentage of wrong calls per site on chromosomes 10-17, total and separated by error mode for LAI ran on the model reflecting a pulse of admixture at 12 generations ago with 15% AMR/ 60% EUR/ 25% AFR proportions.

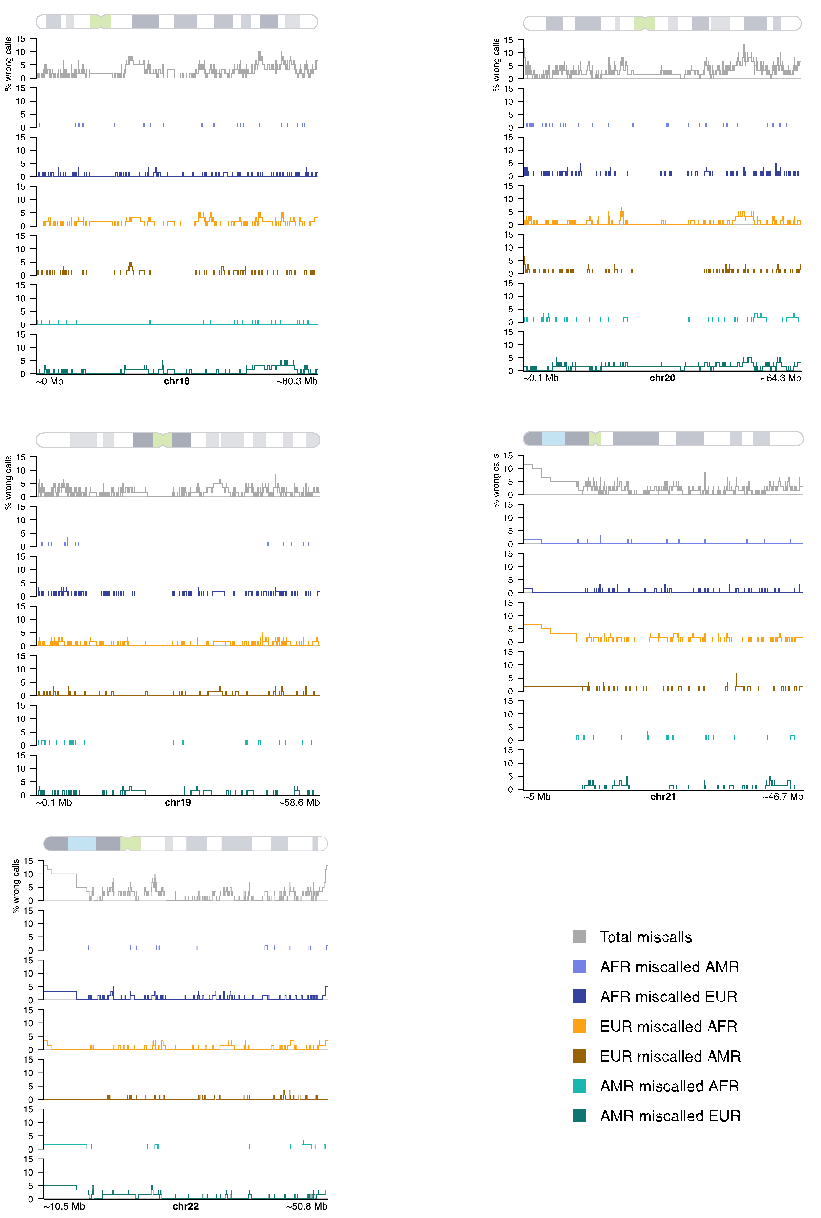

**Supplementary Figure 5:** Percentage of wrong calls per site on chromosomes 18-22, total and separated by error mode for LAI ran on the model reflecting a pulse of admixture at 12 generations ago with 15% AMR/ 60% EUR/ 25% AFR proportions.

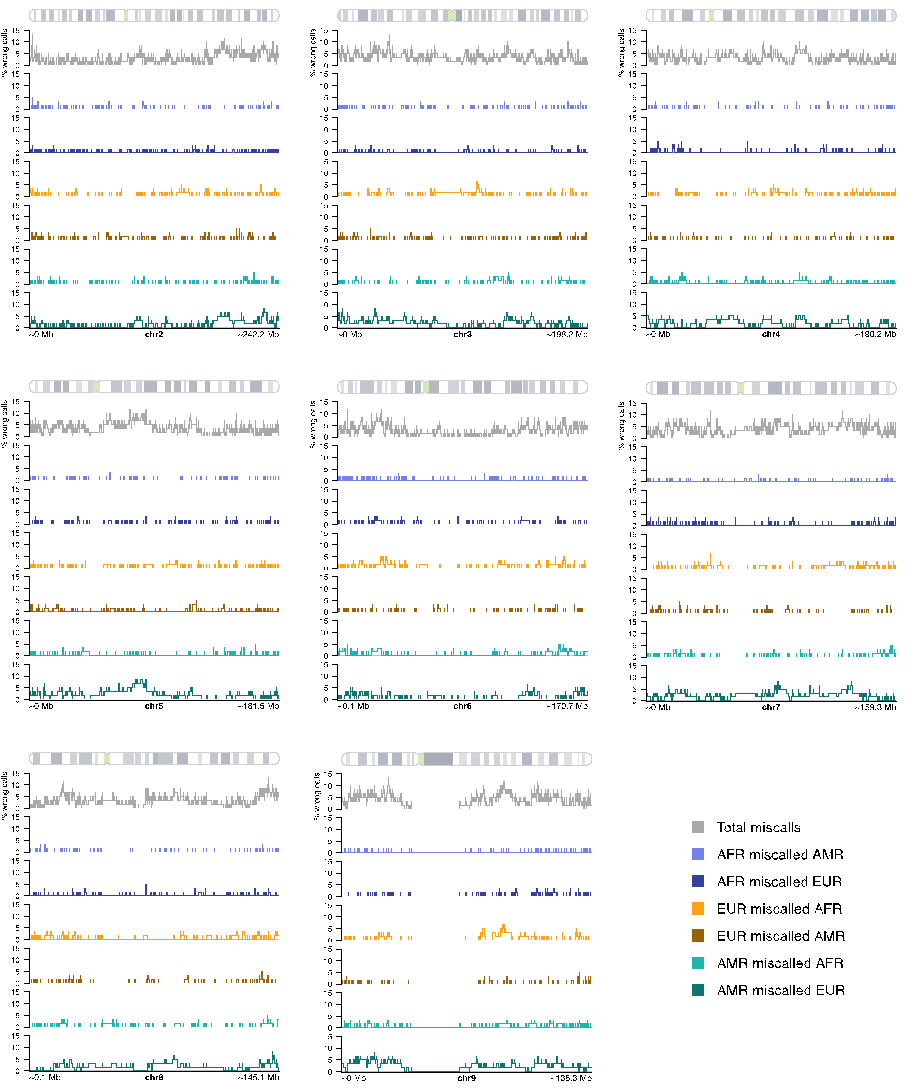

**Supplementary Figure 6:** Percentage of wrong calls per site on chromosomes 2-9, total and separated by error mode for LAI ran on the model reflecting a pulse of admixture at 12 generations ago with 33% AMR/ 33% EUR/ 34% AFR proportions.

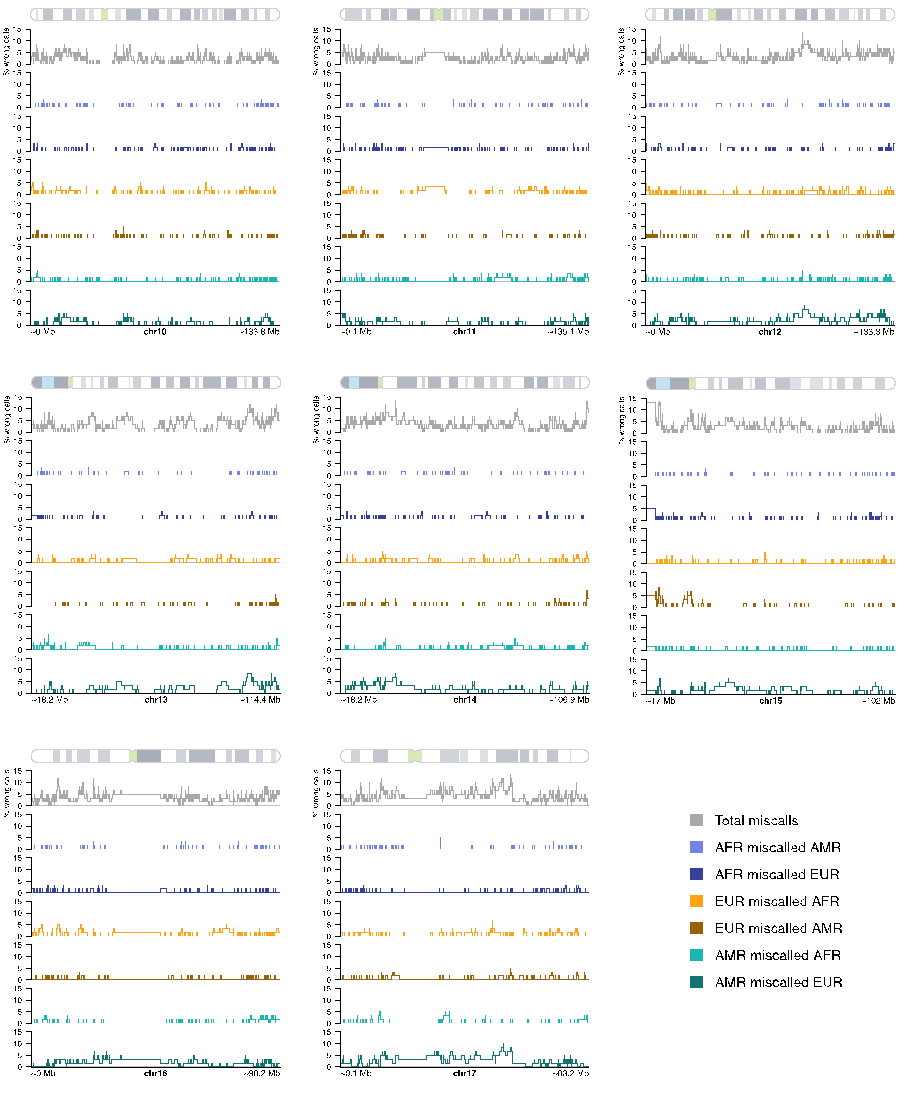

**Supplementary figure 7:** Percentage of wrong calls per site on chromosomes 10-17, total and separated by error mode for LAI ran on the model reflecting a pulse of admixture at 12 generations ago with 33% AMR/ 33% EUR/ 34% AFR proportions.

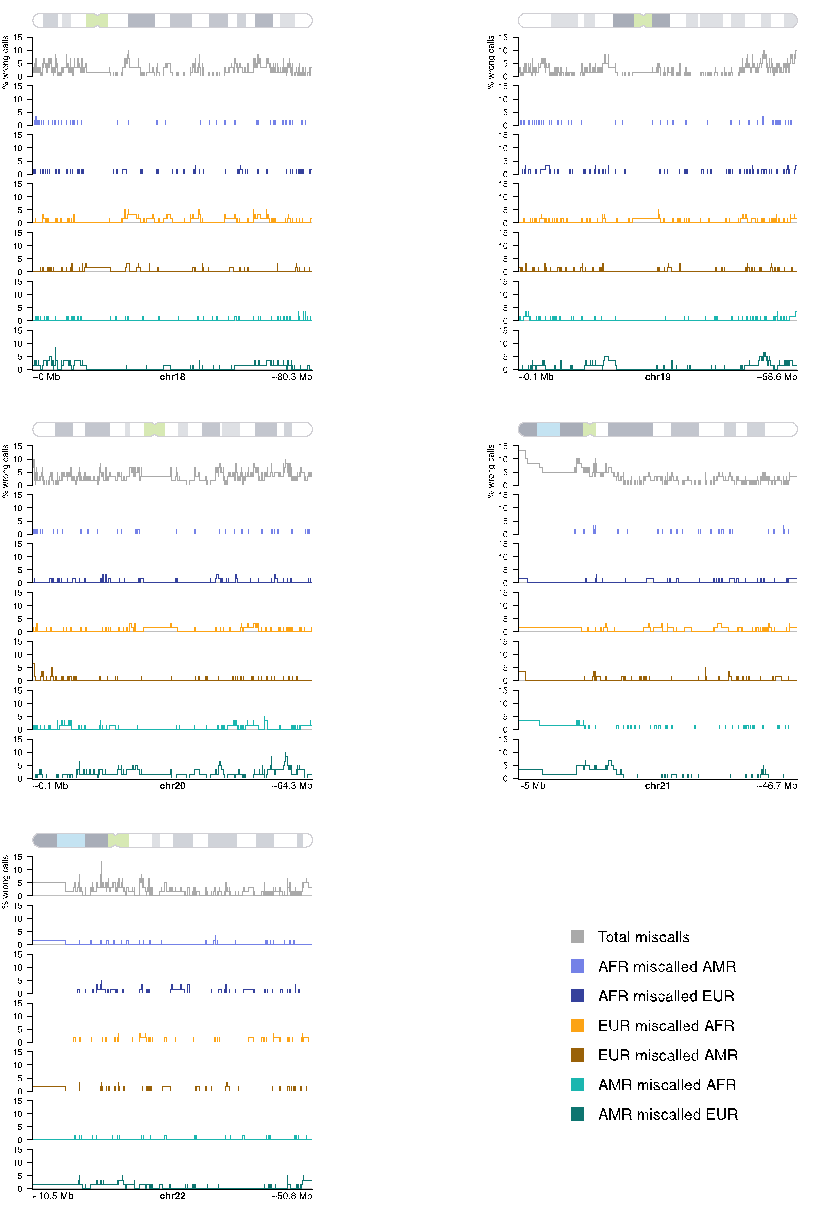

**Supplementary figure 8:** Percentage of wrong calls per site on chromosomes 18-22, total and separated by error mode for LAI ran on the model reflecting a pulse of admixture at 12 generations ago with 33% AMR/ 33% EUR/ 34% AFR proportions.

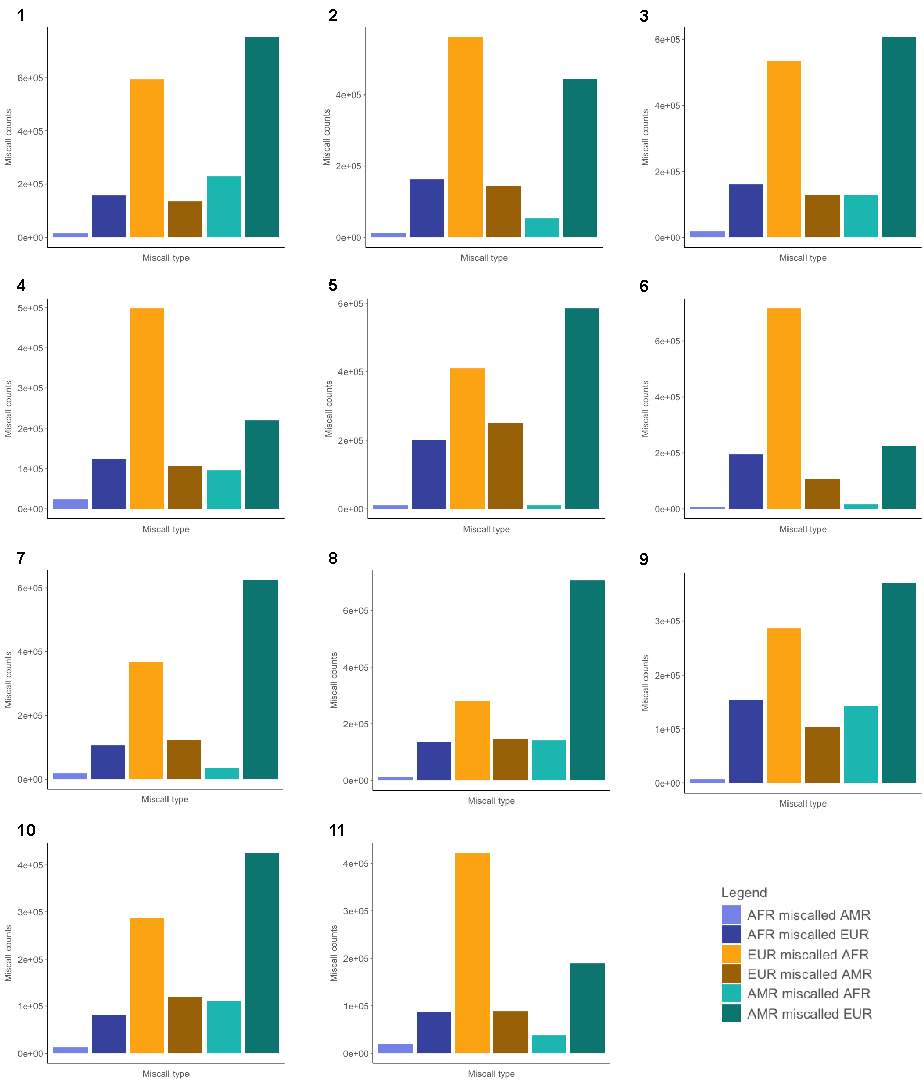

**Supplementary Figure 9:** Miscall counts separated by error mode on chromosomes 1-11 for LAI ran on the ‘average 3-way’ model reflecting a pulse of admixture at 12 generations ago with 15% AMR/ 60% EUR/ 25% AFR proportions (average three-way proportions model).

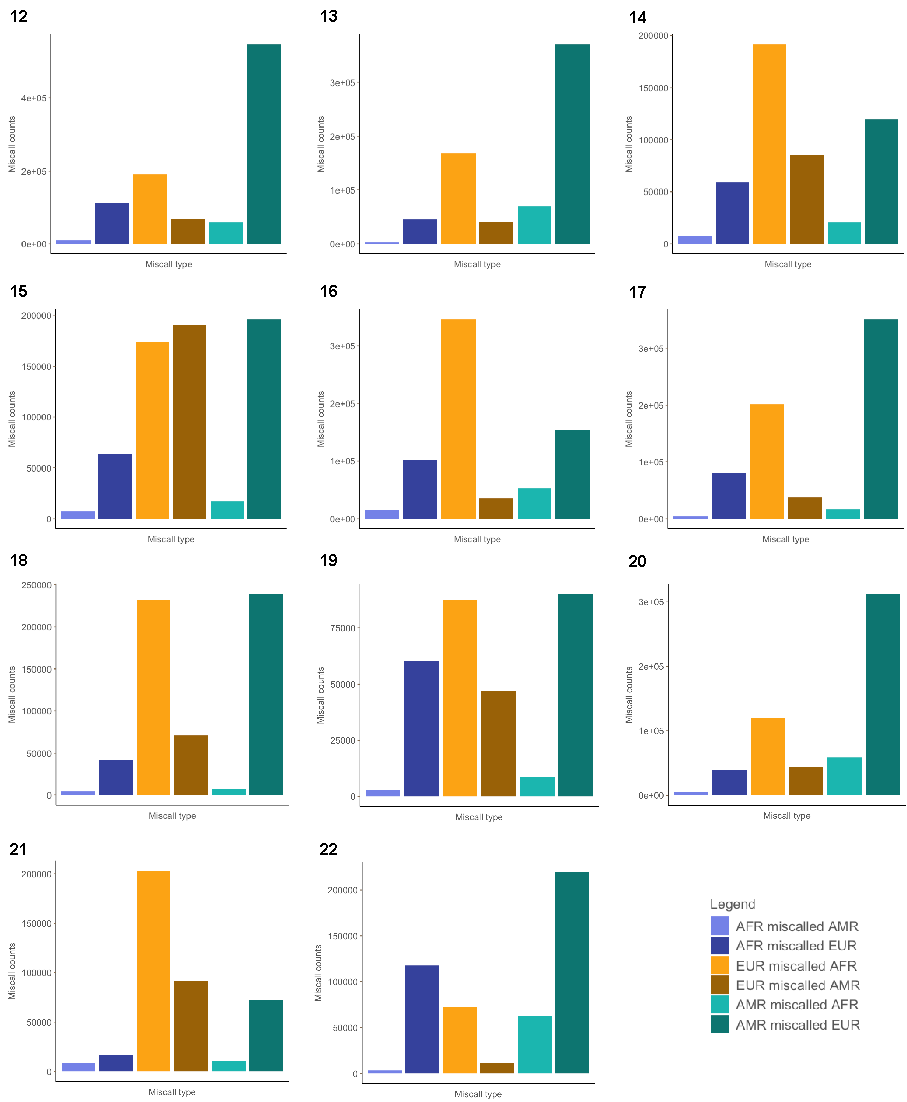

**Supplementary Figure 10:** Miscall counts separated by error mode on chromosomes 12-22 for LAI ran on the model reflecting a pulse of admixture at 12 generations ago with 15% AMR/ 60% EUR/ 25% AFR proportions (average three-way proportions model).

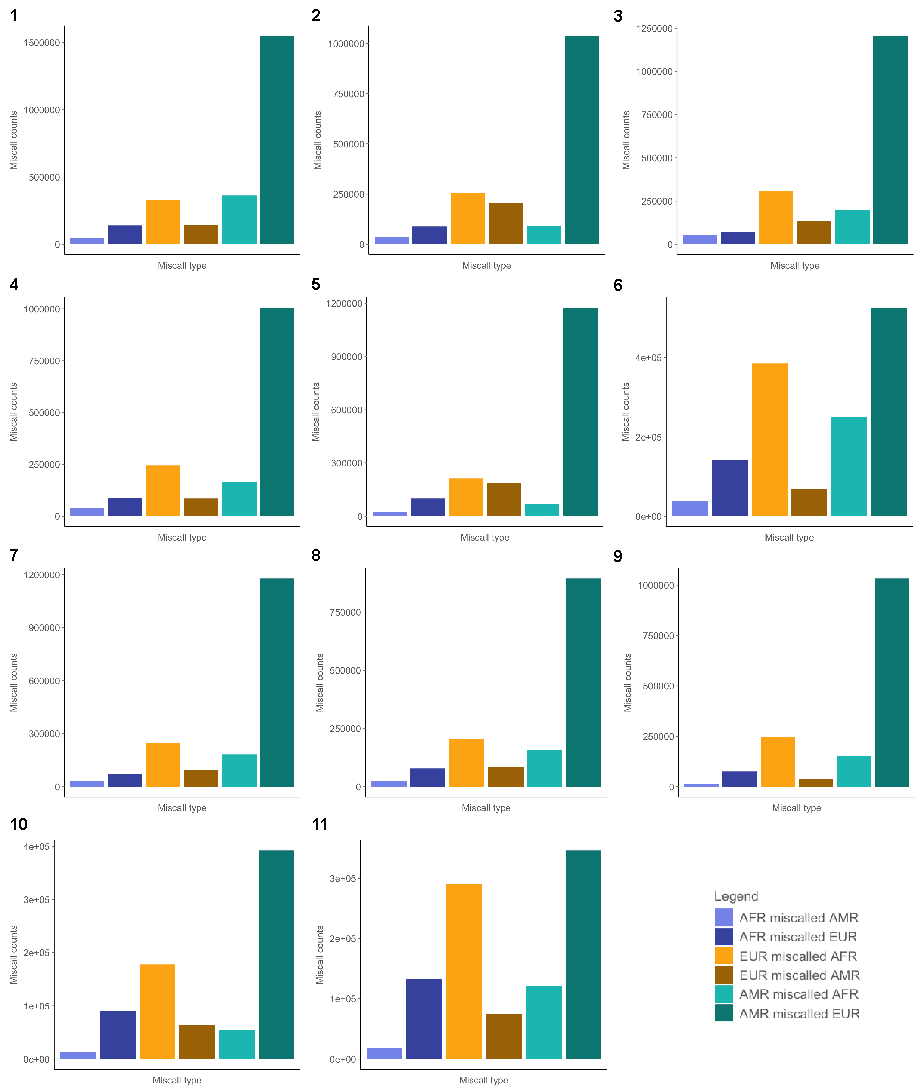

**Supplementary Figure 11:** Miscall counts separated by error mode in chromosomes 1-11 for LAI ran on the model reflecting a pulse of admixture at 12 generations ago with 33% AMR/ 33% EUR/ 34% AFR proportions (even three-way proportions model).

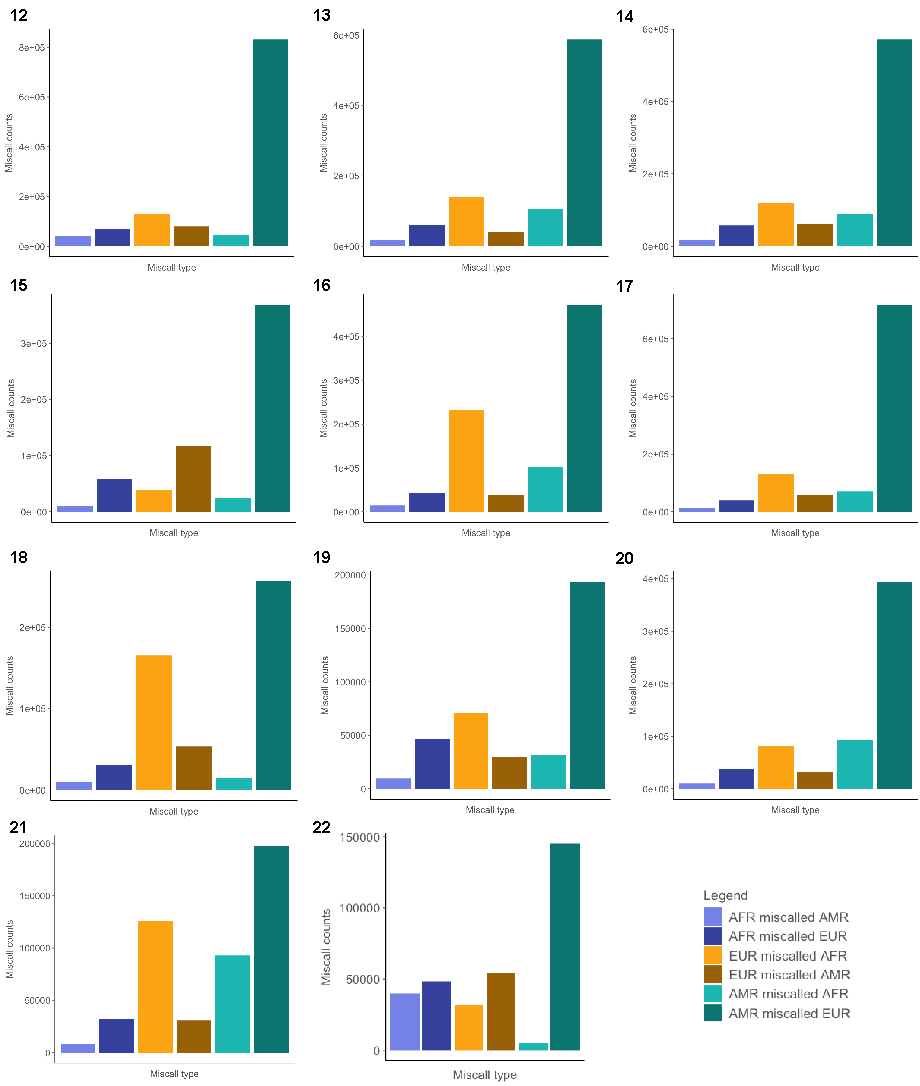

**Supplementary Figure 12:** Miscall counts separated by error mode in chromosomes 12-22 for LAI ran on the model reflecting a pulse of admixture at 12 generations ago with 33% AMR/ 33% EUR/ 34% AFR proportions (even three-way proportions model).
